# Avoiding misleading estimates using mtDNA heteroplasmy statistics to study bottleneck size and selection

**DOI:** 10.1101/2022.09.06.506828

**Authors:** Konstantinos Giannakis, Amanda K. Broz, Daniel B. Sloan, Iain G. Johnston

## Abstract

Mitochondrial DNA (mtDNA) heteroplasmy samples can shed light on vital developmental and genetic processes shaping mtDNA populations. The sample mean and sample variance of a set of heteroplasmy observations are typically used both to estimate bottleneck sizes and to perform fits to the theoretical “Kimura” distribution in seeking evidence for mtDNA selection. However, each of these applications raises problems. Sample statistics do not generally provide optimal fits to the Kimura distribution and so can give misleading results in hypothesis testing, including false positive signals of selection. Using sample variance can give misleading results for bottleneck size estimates, particularly for small samples. These issues can and do lead to false positive results for mtDNA mechanisms – all published experimental datasets we re-analysed, reported as displaying departures from the Kimura model, do not in fact give evidence for such departures. Here we outline a maximum likelihood approach that is simple to implement computationally and addresses all of these issues. We advocate the use of maximum likelihood fits and explicit hypothesis tests, not fits and Kolmogorov-Smirnov tests via summary statistics, for ongoing work with mtDNA heteroplasmy.

## Introduction

Mitochondrial DNA (mtDNA) encodes vital energetic apparatus. Maintaining mtDNA integrity in the face of mutational pressure is a universal challenge for eukaryotes. Several taxa attempt this maintenance in part through segregation of mtDNA damage (and, in photosynthetic taxa, plastid or ptDNA damage) – inducing cell-to-cell variance in germline mutant load so that, while some germ cells inherit high mutant loads, some inherit lower loads and can give rise to viable offspring. This process is often pictured as a genetic “bottleneck”, where a reduced effective population size increases cell-to-cell variability (Johnston, 2019; Stewart and Chinnery, 2015).

Quantifying this increase of variability (the “size” of the bottleneck) with samples of observed genetic data is of interest and importance from fundamental biology to clinical planning. It is common to estimate bottleneck size using the sample mean *ħ* = 1/*n* Σ *h* and sample variance *s*^2^ = 1/(*n*-1) Σ (*h*-*ħ*)^2^ of a set of heteroplasmy measurements. More recently, fits of such measurements to a theoretically predicted heteroplasmy distribution under neutral drift (Kimura, 1955) have been used to explore evidence for mtDNA selection (Wonnapinij *et al*., 2008, 2010; Freyer *et al*. 2012; Zhang *et al*. 2021; Monnot *et al*. 2011; Jokinen *et al*. 2016; Otten *et al*. 2018) and to model mtDNA inheritance (Samuels *et al*. 2013). Within the mtDNA field, this neutral-drift distribution is commonly referred to as the “Kimura” distribution, although Kimura actually derived several other distributions describing the role of selective differences and other features (Kimura, 1954; see below). These fits are also typically performed using the sample mean and sample variance for a set of measurements, in a method outlined by Wonnapinij *et al*. (2008; 2010) and referred to as the WCS-K approach after those authors and Kimura (Wallace and Chalkia, 2013). The interpretation here is that if the Kimura distribution – which assumes no selective differences – does not fit the data well, there may be some support for selective differences in the system of interest.

Specifically, bottleneck size is typically estimated using

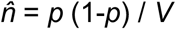

where *p* and *V* are sample statistics, calculated from observations in different ways in different projects. Most commonly, given a sample of heteroplasmy values, the sample mean *ħ* is used for *p* (thus assuming an absence of selection (Johnston, 2019)) and the sample variance *s*^2^ for *V*. In other cases, an earlier or reference heteroplasmy measurement may be used for *p*; the population variance expression is sometimes used for *V*. It should be remembered that all of these quantities are estimators, based on a sample, for the population relationship *n* = *p*_0_ (1-*p*_0_) / *σ*^2^, which in turn comes from a binomial model where the population variance is given in terms of the parameters 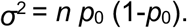

The Kimura distribution as used by the WCS-K approach takes two parameters *p* (initial heteroplasmy) and *b* (describing the extent of drift), both between 0 and 1. A value of *b* close to 1 corresponds to little drift and variance; lower values correspond to more drift and higher variance (*b* is connected to bottleneck size *n* via *n* = 1/(1-*b*)). The population mean and variance are

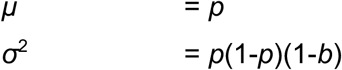

Typically, a fit to observed heteroplasmy samples is performed by calculating the sample mean and sample variance of the sample, asserting that these are equal to the population mean *μ* and population variance *σ*^2^ above, and using these relationships to provide estimates for the parameters *p* and *b* (Wonnapinij *et al*., 2008; 2010). If the equality assertion is instead recognised as an estimation, this is the so-called “method of (central) moments” for fitting a distribution.

### The problems

The problems this article addresses can be demonstrated with four related examples.

*Problem 1. Bottleneck sizes under one*. Consider taking two heteroplasmy measurements in two samples, and observing one homoplasmic wildtype and one homoplasmic mutant *<u>h</u>* = (0, 1). The sample variance is:

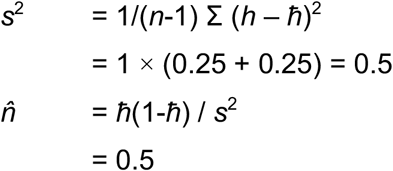

Leaving us with a bottleneck size of 0.5 segregating units. Such a value is nonsensical, because we cannot segregate less than a single unit of information through a bottleneck (even a theoretical, binomial one). If we enforce a particular initial heteroplasmy measurement instead of using the sample mean, the problem remains and can be amplified (see Appendix). This is not an unrealistically contrived case: for example, in recent work revealing segregation dynamics in *Arabidopsis*, Broz *et al*. (2022) found several instances of such bi-homoplasmic sets for mtDNA (and ptDNA), giving bottleneck size estimates under one.

*Problem 2. Uninterpretable parameter fits to, and uncertainty from, the Kimura distribution*. We can immediately see that fitting the Kimura distribution based on matching sample statistics is not generally correct, by considering our *<u>h</u>* = (0, 1) sample. We would use:

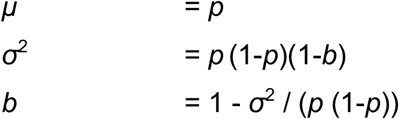

so for our (0, 1) sample, if we use the sample mean *ħ* = 0.5 as an estimate for population mean p and the sample variance *s*^2^ = 0.5 as an estimate for population variance *σ*^2^, we estimate the nonsensical *b* = 1 – 0.5 / 0.25 = −1, which cannot then be taken forward into an uncertainty estimate (*b* must take values between 0 and 1). We have also identified more general issues with our ability to calculate uncertainty on our bottleneck estimate using a method-of-moments fit to the Kimura distribution (see below).

*Problem 3. False positive signals of selection (or other processes) via Kimura fit*. Consider the dataset *<u>h</u>* = (0, 0, 0, 0, 0.9, 0.9, 0.9, 0.9). As before, using the WCS-K approach, we fit the Kimura distribution using summary statistics *p* = *ħ* = 0.45 and *b* = *s*^2^ = 0.0649. A Monte Carlo implementation of the Kolmogorov-Smirnov (KS) test (Wonnapinij *et al*., 2008) gives p-values between 0.035 and 0.04, suggesting a significant departure from the theoretical model. Such a departure could be interpreted as a signature of selection.

But now consider a different Kimura distribution, with *p* = 0.392 and *b* = 0.252 (the provenance of these parameters will be explained below). Now the KS test gives p-values between 0.53 and 0.55 (see Fig. 1 below). So there is no reason to reject the null hypothesis that these observations were drawn from a Kimura distribution – removing our reason to suggest that selection may be acting. Because we previously used an (inappropriate) fit based on summary statistics, we chose the wrong Kimura distribution with which to compare our data, and obtained a false positive. Here, we have chosen a contrived dataset to simply illustrate this point, but the issue is systemic: upon reanalysing a range of real datasets for which significant departures were reported (see below), we have not identified any cases that in fact departed significantly from the Kimura distribution at a threshold of p < 0.05.

**Figure 1.**
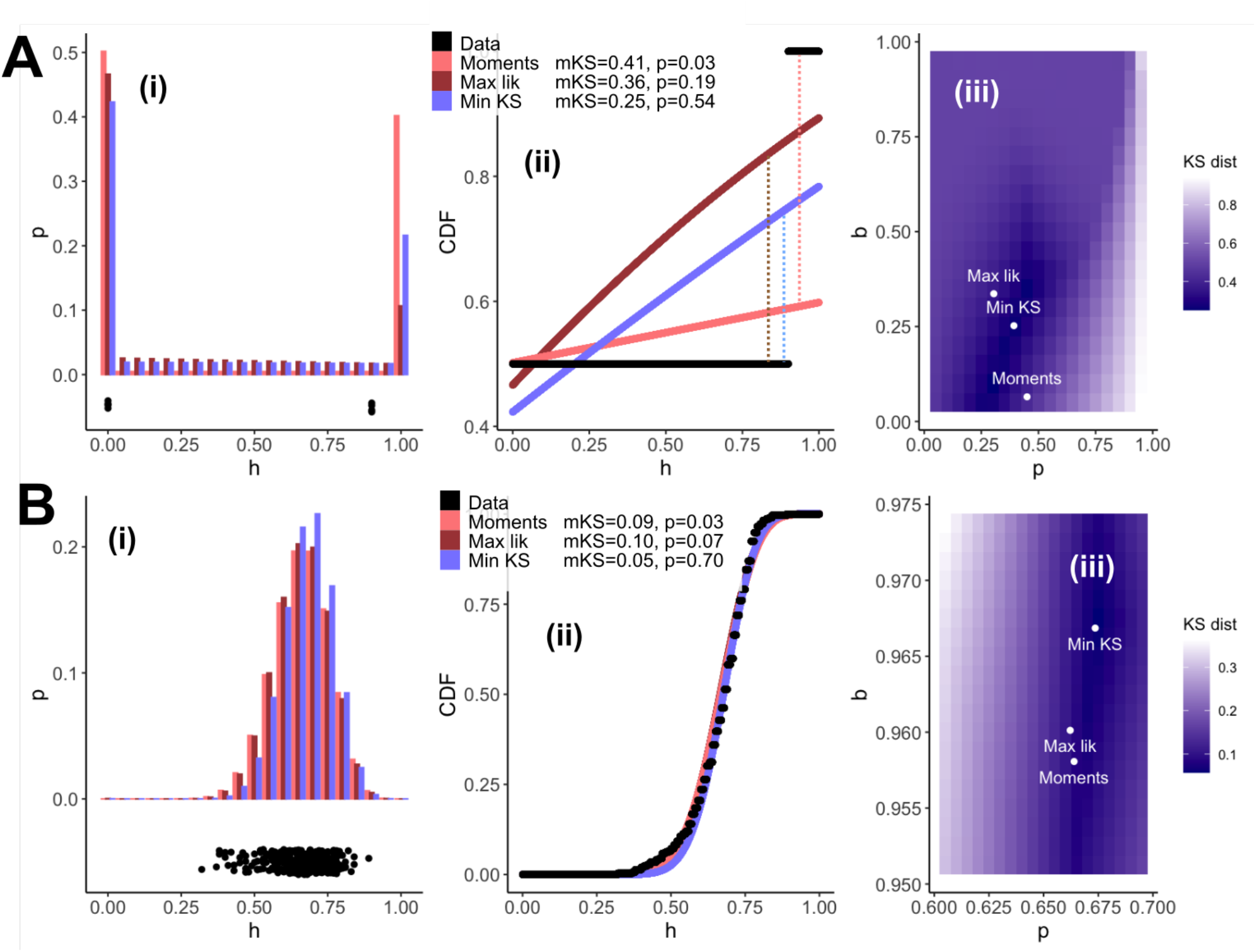
Different estimators for the Kimura distribution give dramatic differences in hypothesis testing. Fits via the (commonly implemented) method of moments, maximum likelihood, and minimum Kolmogorov-Smirnov distance to example datasets. (A) Synthetic data; (B) data from Freyer *et al*. (2012) (specifically, the concatenated set of observations for mother heteroplasmies > 0.7). (i) Probability distributions for each fit; (ii) comparison of cumulative distribution functions (CDFs) from the data and for each fit; (iii) the KS distance between empirical observations and theoretical distribution with parameters of the Kimura distribution. mKS, KS distance for each method of fitting; p-values are from the Monte Carlo KS test in WCS-K. The maximum distance between CDFs of the data and a fitted distribution gives the KS distance (illustrated by dotted lines in (A)(ii)); the method of moments (and maximum likelihood) give KS distances rather higher than the minimum KS distance approaches, with correspondingly low (and false positive) p-values in hypothesis testing. Even the subtle differences in parameterisation for (B) give dramatically different p-values in the Kolmogorov-Smirnov test.

*Problem 4. Hypothesis testing with bottleneck sizes*. Sample variances are not drawn from a normal distribution. Even when the population from which samples are taken is normal, the sample variance follows a chi-squared distribution; when the population distribution is not normal (as for heteroplasmy measurements), the sample variance distribution is in general not known. We cannot then use tests that invoke a normality assumption (like the t-test) to compare sample variances or the bottleneck size estimates that are derived from them. Must we use low-powered non-parameteric approaches instead?

## Results

### A parametric solution – maximum likelihood fitting

How can we estimate a meaningful bottleneck size and derive sensible confidence intervals in the challenging cases above? Fitting heteroplasmy measurements to the Kimura distribution can actually answer all of these questions. However, the above examples make it clear that we cannot in general perform a simple matching of distribution parameters to summary statistics (choosing the parameters, *p* and *b*, that give a distribution with the same mean and variance as the sample). This method-of-moments will only find the parameters that are most compatible with the observations in the case of infinite sample size, where the sample mean and variance converge to the population mean and variance.

When individual heteroplasmy measurements are available (sometimes only summary statistics are reported), and there is reason to believe the Kimura model, a more appropriate approach is instead to identify the maximum likelihood parameters given the full set of measurements (as used previously in at least one study (Otten *et al*., 2018)). The maximum likelihood parameters for a statistical model are those that give the highest joint probability of observing our measurements under that model. Although these do correspond to the sample mean and sample variance in the case of a normal distribution model, for other distributions (including the Kimura distribution) this is not generally the case (as in the example above). Instead we have to find the parameters with the highest associated likelihood. The Kimura distribution Kimura(*h* | *p, b*) gives the probability density for a given heteroplasmy observation *h*. We write

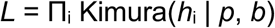

and seek the *p, b* combination that maximises *L* for a given *h*. If we wish to enforce a particular *p* – for example, if we have a reliable initial heteroplasmy measurement – we can instead perform the search only over *b*. Both *p* and *b* are confined in [0,1] here, respecting the constraints of the system.

This maximum likelihood process will identify the *b* (and *p*, if required) that is most likely given the set of observations. We can also derive confidence intervals on this estimate using Fisher information or bootstrapping (see Appendix), obtaining, for example, an estimate for *b* and its 95% confidence intervals. These can then be interpreted as bottleneck size *n* via *n* = 1/(1-*b*).

To reiterate, this process is not the same as fitting a Kimura distribution based on summary statistics, as is often used. In that case we are losing information about the distribution of heteroplasmy samples and will not in general identify the maximum likelihood parameterisation. How does this approach deal with our examples above? First consider the *<u>h</u>* = (0, 1) case. A maximum likelihood fit gives us *p* = 0.5 (95% c.i.s 0.058-0.941), and a point estimate of *b* ≃ 0 (and corresponding *n* = 1) – all readily interpretable as well-behaved parameters of the Kimura distribution. Confidence intervals on *n* can be estimated via taking likelihood derivatives or resampling, the latter of which is better behaved for this small, extreme dataset (see Appendix), and gives *n* between 1 and 13.6 with 95% confidence.

Applied to the larger sets of model observations, the maximum likelihood approach readily identifies the most likely model parameters given observations (Fig. 2). In these cases the confidence intervals on *n* can readily be computed using likelihood derivatives, immediately giving an interpretable uncertainty on bottleneck size.

**Figure 2.**
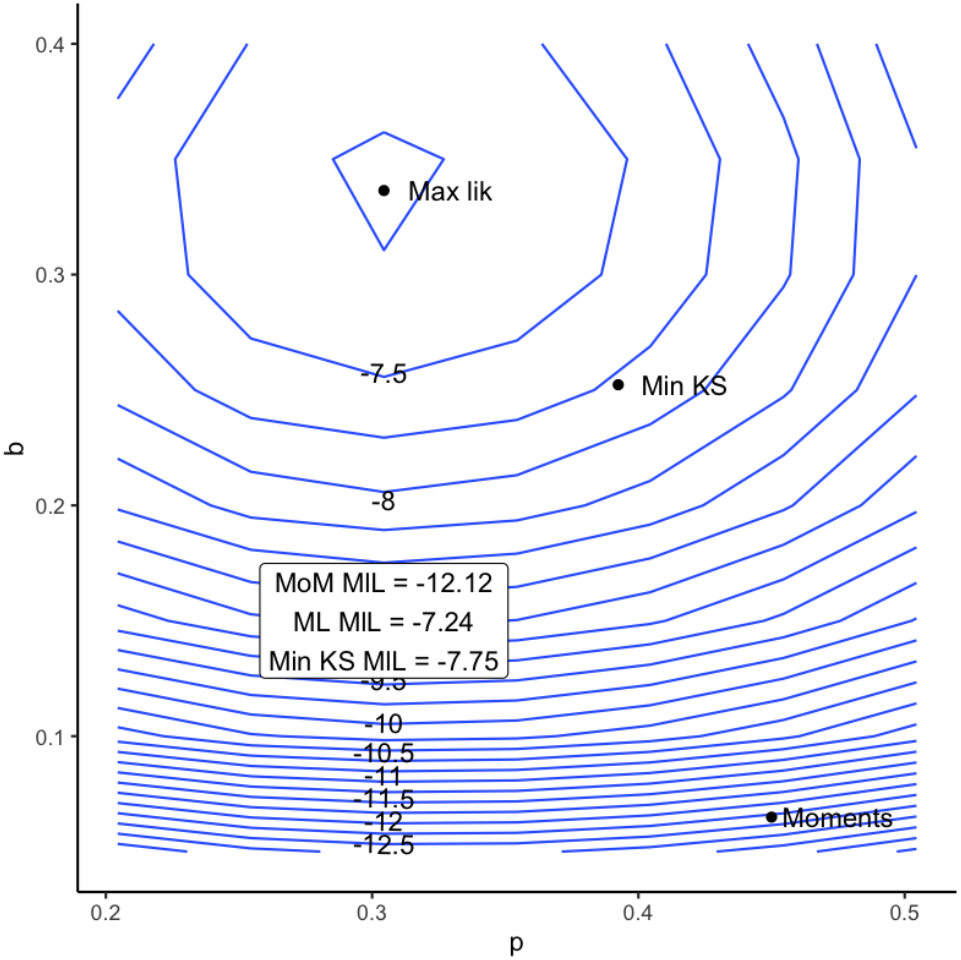
Likelihoods for different model fits. Likelihood surface for the synthetic dataset in Fig. 1A, with the parameterisation from different model fits illustrated. The maximum likelihood fit by construction identifies the parameterisation with the highest likelihood; the method of moments and minimum KS distance fits identify parameterisations some distance from this peak.

### Testing fits to the Kimura distribution

Wonnapinij *et al*. (2008) propose using a Monte Carlo method based on the Kolmogorov-Smirnov statistic between the empirical cumulative distribution function of a heteroplasmy sample and an ensemble of samples generated from a fitted Kimura distribution. However, this fitting is typically carried out using the method of moments, which is not guaranteed to give a distribution that will generate samples with the lowest KS distance from the data. The maximum likelihood approach above does not solve this problem: the maximum likelihood parameterisation of the Kimura distribution is also not guaranteed to minimise KS distance from the data. In general, fitting by moments, likelihood, and goodness-of-fit (including KS distance) will give different estimators for finite samples (Figs. 1, 2).

It is possible to find the parameterisation of the Kimura distribution that *is* guaranteed to give the minimum KS distance from a given dataset, by minimising the KS distance across parameter values (Fig. 1; see Appendix). This parameterisation can yield much lower KS distances than the two other estimators. In all cases we studied, this means that the p-value computed from the Monte Carlo approach does not pass a significance threshold, even when the other parameterisations (less suited to minimising KS distance) suggested a significant departure. Some examples are shown in Fig. 1; this is the approach used to identify the alternative parameterisation in the motivating example problem above.

In other words, the fact that the Monte Carlo approach gives a low p-value when used with a distribution fitted via moments does not mean that the Kimura distribution is incompatible with observations. It is very possible that a different fit, minimising KS distance instead of matching moments, will give a p-value that does not pass a significance threshold – so the hypothesis that a Kimura distribution (just not the moment-fitted one) generated the data cannot be discarded. We found this to be the case in every dataset we investigated, including several where a significant departure from the Kimura distribution was previously reported (Wonnapinij *et al*., 2008; Freyer *et al*. 2012) (Table 1). Other studies using the Kimura fit did not report significant departures from the Kimura distribution (Zhang *et al*. 2021; Monnot *et al*. 2011; Jokinen *et al*. 2016); in these cases the minimum KS distance fit will simply yield even higher p-values (as in Fig. 1 and Table 1).

**Table 1.**
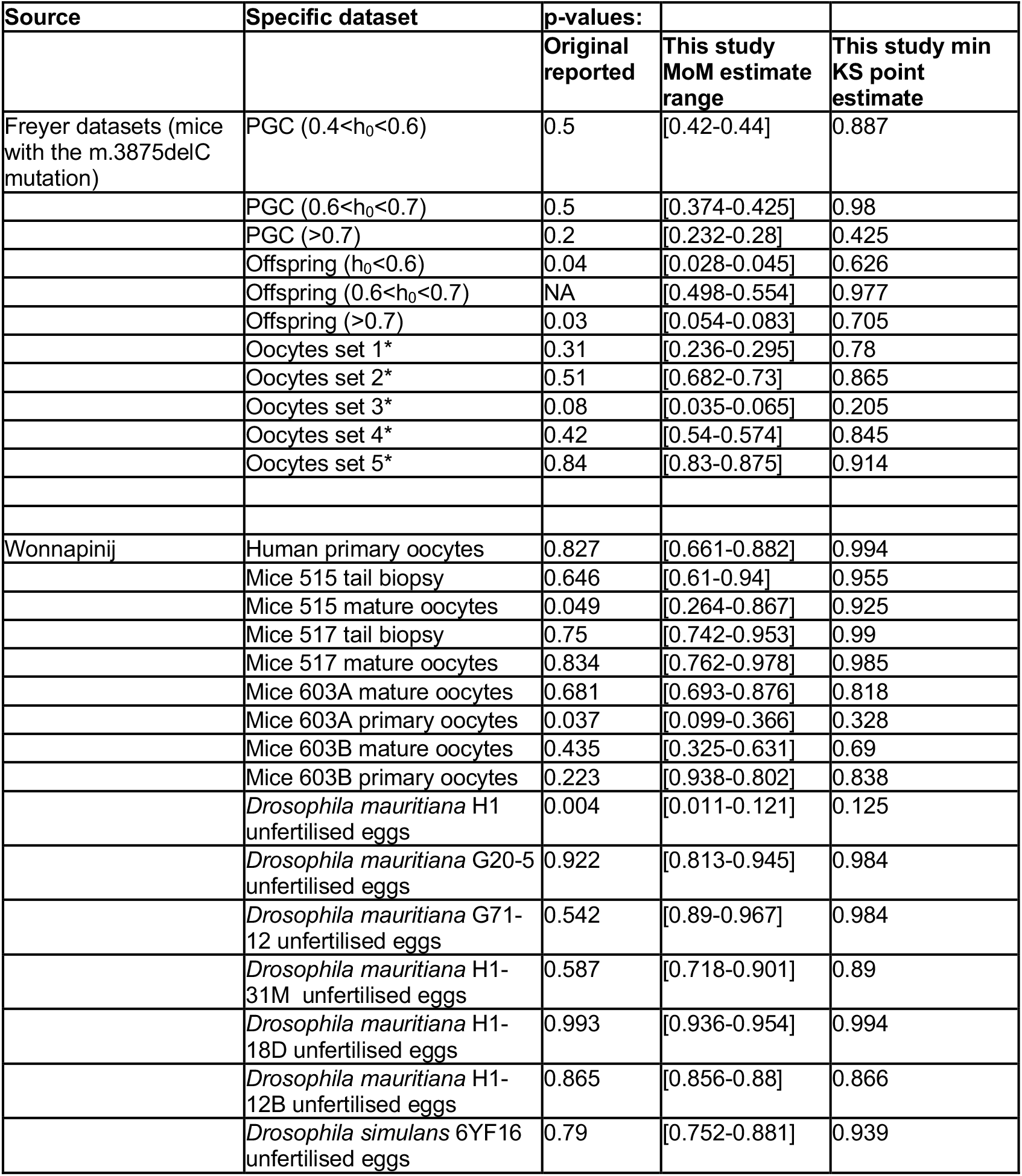
Appropriate parameter fits to the Kimura distribution remove statistically significant signals of selection. Comparison of reported p-values (from method-of-moments fits) with the range of p-values from method-of-moment fits in this study, and p-values from minimum KS distance fits. Data are from several original sources where significant departures from the Kimura distribution were reported (Wonnapinij *et al*., 2008; Freyer *et al*. 2012). In several cases, our method-of-moments p-value estimate range departs from that originally reported, illustrating the sensitivity of KS p-values on the specific values used (see Discussion). See Methods for details of how data were collected and parsed. * The ordering of these experiments was not specified in the original publication; we have ordered them to align as much as possible with our estimated MoM p-values.

The fact that judicious parameterisations of the Kimura distribution can fit this wide range of heteroplasmy distributions underlines that the Kimura distribution is remarkably flexible, and capable of capturing a wide range of heteroplasmy structures (including all those observed to date, to our knowledge). Despite this flexibility, it is possible to construct a dataset that cannot be well fit (in the KS sense) by any parameterisation of the Kimura distribution. An example is given by expanding the above (0, 0.9) example so that there are 50 observations of each value (p = 0.01 from Monte Carlo KS test using minimum KS distance fit, Supplementary Figure 1). This structure is challenging to fit because, on one hand, the many zeroes suggest either a low *p* or very high segregation, and the many 0.9s suggest neither can be true. However, this and comparable examples are far removed from any real-world observations of which the authors are aware.

For clarity, it helps to specify the hypotheses that these various approaches test. The original approach tests the hypothesis that a specific Kimura distribution, parameterised by the method of moments, commonly generates samples with a higher KS distance from the distribution than the data’s KS distance. Because that particular parameterisation is not guaranteed to reflect the true distribution, this test is hard to interpret. The proposed approach tests the hypothesis that any Kimura distribution commonly generates samples with a higher KS distance from the distribution than the data’s KS distance. This test is more (but not fully) aligned with the scientific question of whether the data may be generated given the assumptions underlying the Kimura model.

The question of whether samples are unlikely to be drawn from a given family of distributions is complicated. While results like variants of the Anderson-Darling test exist for many distributions (Stephens, 2017), to our knowledge these results do not exist for the Kimura distribution. We suggest that such tests are not yet developed enough to provide scientific insight, and instead advocate the likelihood-based testing of alternative hypotheses as in Otten *et al*. (2018).

### Non-parametric solutions I – h-statistics

In some cases this maximum likelihood approach may be impossible (if we do not have access to individual measurements) or undesirable (if we do not believe that the Kimura distribution, or any other parametric model, is a good description of the system). In such cases we may be forced to use a non-parametric approach to estimate bottleneck size. Here, there is no way of avoiding some of the issues above, as without a model we cannot naturally enforce scientific constraints on the values involved. With this caveat, sample statistics can be used to provide an estimator of the uncertainty in sample variance (Wonnapinij *et al*., 2010). However, as shown in our introductory problems, several issues can arise with this approach and require careful interpretation; we also believe that the expressions in Wonnapinij *et al*. (2010) need some adjustment to be generally applicable.

The variance of the sample variance *s*^2^ is

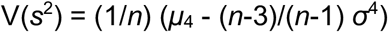

which requires two population quantities, the variance *σ*^2^ and the fourth central moment *μ*_4_, to be estimated from a sample of data.

Wonnapinij *et al*. (2010) quote a result for a quantity *D*_4_, which is proposed as an unbiased estimator of the fourth central moment of a distribution

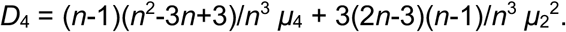

However, we cannot find justification for this estimator. In the cited source (Dodge and Rousson, 1999), the left hand side of this equation (Wonnapinij’s *D*_4_) is not presented as an estimator of *μ*_4_, but is the expected value of the sample moment *m*_4_. The reference states that the expected value of that sample moment is the expression on the right hand side, not that this is an estimator for the population *μ*_4_. Indeed, *μ*_4_ (the quantity to be estimated) itself appears on the right hand side. In tandem, several other key expressions in the WCS-K approach, including for the normal and Kimura special cases, involve population quantities *σ*^2^, *p, b*, and even *μ*_4_ itself, which are not directly accessible from a sample.

Happily, all this is resolved by the existence of a unique unbiased symmetric estimator for *μ*_4_ in terms of sample moments, which is the corresponding h-statistic (Rose and Smith, 2002; Halmos, 1946; the h label here and *h* for heteroplasmy are a coincidence):

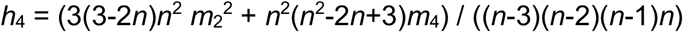

where

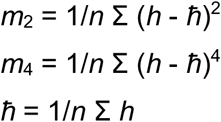

(note that these expressions are all functions of the data sample alone, not population quantities as in the WCS-K expressions). Here we can immediately take our *n* measurements of *h*, compute sample mean *ħ* and sample moments *m*_2_ and *m*_4_, and obtain our *h*_4_ estimate of *μ*_4_ for further use.

The expression in Wonnapinij *et al*. (2010) for *μ*_4_ in the Kimura distribution is correct (the algebra required is in the Appendix) but, as before, this is a population quantity and we cannot in general simply plug in a set of estimators and obtain an unbiased *μ*_4_ estimate. Using the h-statistic immediately resolves this issue.

We therefore propose the following estimator for the variance of the sample variance, based directly on sample statistics from the data

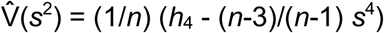

Once this variance estimate has been obtained, its interpretation as confidence intervals requires a parametric choice (for example, writing +-1.96 s.e. invokes a normal assumption). To avoid these issues we can use a resampling approach.

### Non-parametric solutions II – Resampling to estimate variance uncertainty

An alternative approach is possible without relying on a parametric model, particular estimators, and without requiring a parametric choice at any point in the analysis. The bootstrap and jackknife are two algorithms from applied statistics that allow very general estimation of uncertainty on any statistic of interest, computed by “resampling” the data set (Efron and Tibshirani, 1994). This process involves generating a set of new samples from the original sample, related but different, and computing the statistic of interest for each new sample. This set of computed values estimates the true distribution of the statistic of interest. In the bootstrap, *B* new samples of size *n* are constructed by sampling with replacement from the original sample. In the jackknife, *n* new samples of size *n*-1 are constructed by omitting each element of the original sample in turn. We focus on the bootstrap here for simplicity.

Bootstrap estimates for heteroplasmy variance are then constructed by creating *B* new samples and working out the heteroplasmy variance for each, with the uncertainty in the overall estimate given by the spread of values across this resampled set. There is an important technical point here: resampling with replacement leads to bias in summaries of dispersion (like variance) because the same observation will often be repeated in a resampled set (Wiley, 2001). Several methods have been proposed to correct this bias, described and benchmarked in Nguyen (2018). We adopt the simple approach in Wiley (2001) (generalised in Brennan (2007), and supported by the benchmarking across a wide range of cases) – using a correcting factor of *n/(n-1)* to compensate for the expected bias. The estimate of standard error of heteroplasmy is then

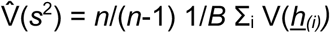

where the sum is over *B* bootstrap resamples, each giving a resampled dataset *<u>h</u>*_*(i)*_.

### A roadmap for heteroplasmy analysis

Taken together, these approaches give us a set of options for the analysis of heteroplasmy data. These options are outlined in the form of a decision tree in Fig. 3.

**Figure 3.**
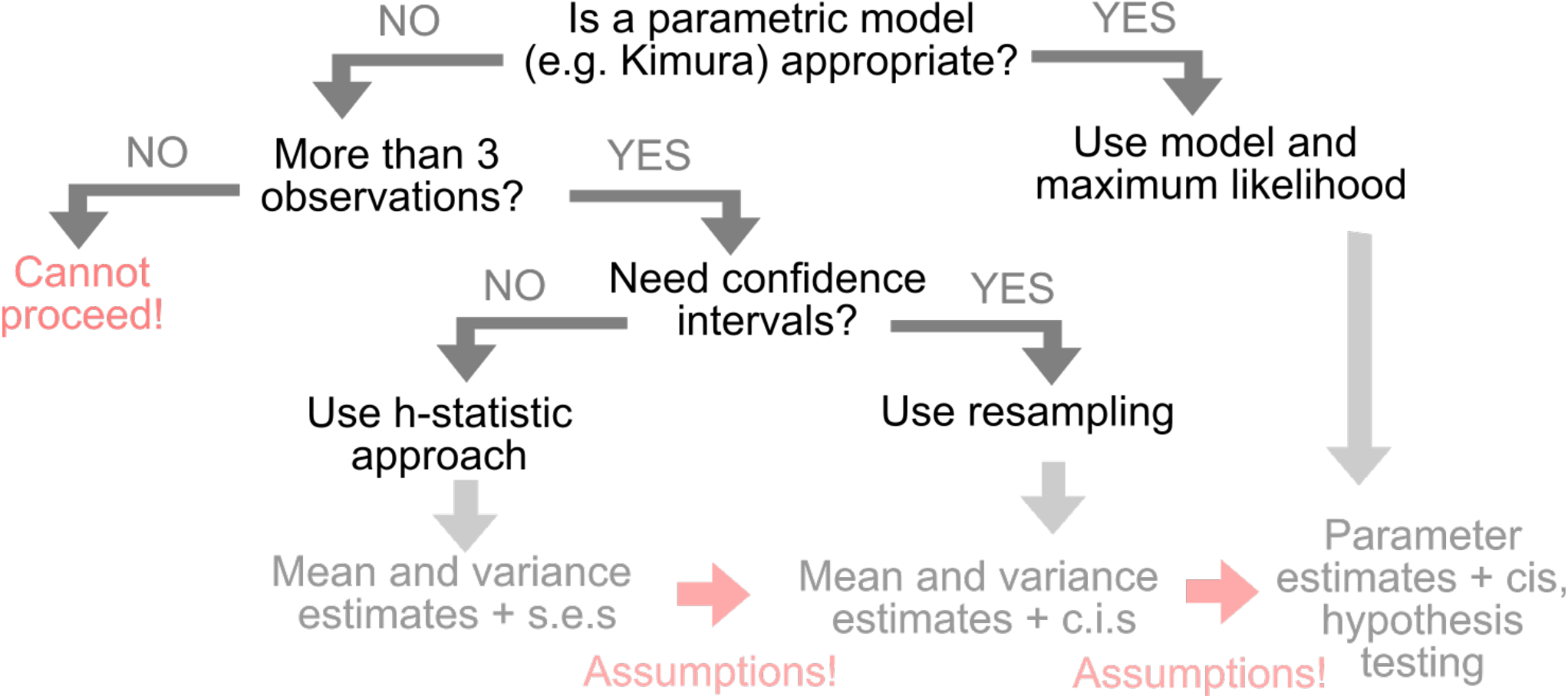
Choosing approaches to analyse heteroplasmy measurements. A decision tree outlining our proposed use of different statistical approaches to analyse heteroplasmy. The “assumptions” involved are (i) a parametric way of interpreting standard errors as confidence intervals (for example, a normal model of +-1.96 s.e.) and (ii) a link between the estimated summary statistics of a dataset and the parameters of a generative model. s.e., standard error; c.i., confidence interval.

In Fig. 4, the different approaches for estimating uncertainty in heteroplasmy variance are illustrated for a selection of the (highly segregated) samples in the plant organelle dataset from Broz *et al*. (2022). In Supplementary Fig. 2, these different approaches are illustrated for a wider range of synthetic and real datasets. Across examples, non-parametric (h-statistic and bootstrap) estimates are typically consistent, and parametric fits to the Kimura distribution give more conservative (larger) estimates for V(h) variance.

**Figure 4.**
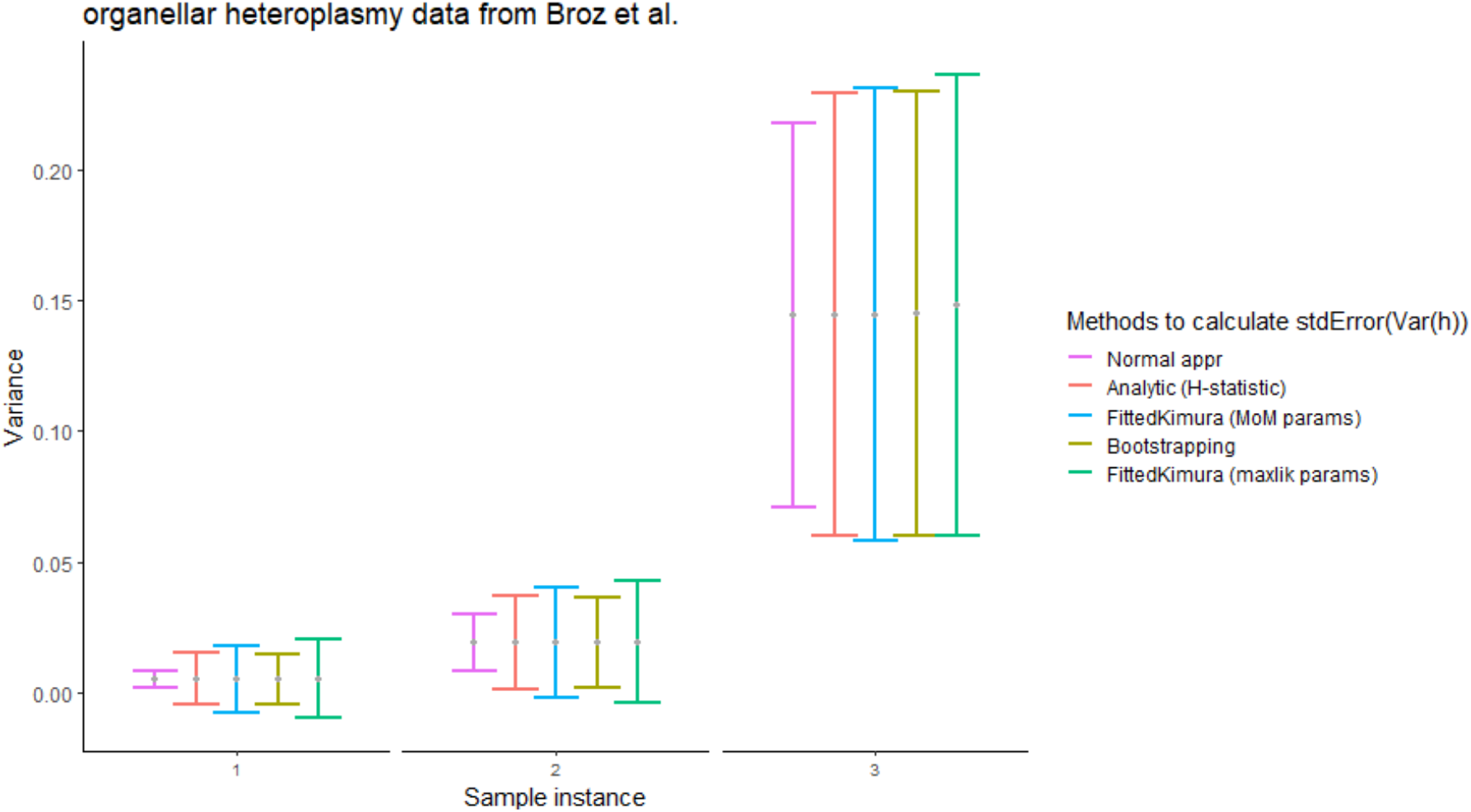
Different ways to estimate the uncertainty of variance in heteroplasmy samples. Variance of a subset of organellar heteroplasmy samples from Broz *et al*. (2022). For comparison, error bars are set to twice the estimated standard error of variance. Non-parametric (h-statistic and bootstrap) approaches give generally consistent estimates; parametric fits to the Kimura distribution are more conservative. Method-of-moments and maximum likelihood fits of Kimura parameters can give different estimates of uncertainty, particularly pronounced in the final set. See Appendix for examples from other experimental studies.

### Bonus results I – Comparing bottleneck sizes

Consider the problem of how to compare bottleneck sizes in different systems. We cannot use a t-test – bottleneck size is the reciprocal of a sample variance and cannot possibly be normally distributed. Non-parametric approaches will lose statistical power. Can we use the above idea to perform well-powered testing of hypotheses about bottleneck size?

Yes, readily. Given the ability to perform maximum likelihood estimates for our parametric models, we can use a likelihood ratio test to compare two models: first, where a different bottleneck applies to different observations, and second, where the same bottleneck describes all observations – following Broz *et al*. (2022). Consider the case where we have two groups, each consisting of several sets of heteroplasmy observations. We are interested in whether the bottleneck size is the same in the two groups. We consider two model structures. First, each set in each is assumed to be drawn from a Kimura distribution, where *p* is unique for each set and *b* is unique for each group. Second, each set is assumed to be drawn from a Kimura distribution, where *p* is unique for each set and the same *b* value applies to both groups. Hence

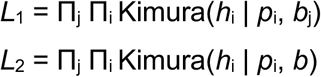

We then maximise *L*_1_ and *L*_2_ over *p*_i_ and *b*_j_ (for model 1) or *b* (model 2). We can then use the likelihood ratio test to investigate support for model 1 (different bottleneck sizes in the two groups) against model 2 (the same bottleneck size in both groups):

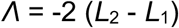

Comparing *Λ* to a chi-squared distribution with one degree of freedom (capturing the one parameter difference between the two models) will then give a p-value against the null hypothesis of no difference in bottleneck size between the groups.

For example, consider two groups: Group 1 with *h*_11_ = (0.2, 0.4), *h*_12_ = (0.6, 0.7) and Group 2 with *h*_21_ = (0.1, 0.7) and *h*_22_ = (0.3, 1). The maximum likelihood process above gives estimates for the bottleneck size of 34 (9.4-130) and 2.3 (1.3-5.8) respectively, and the likelihood ratio test gives a p-value of 0.005 against the null hypothesis of equal bottlenecks.

This p-value may seem surprisingly small given the low sample size in our example. But this is the strength of an appropriately-chosen parametric approach. It is very unlikely that a single Kimura distribution capable of generating the high-variance pairs would also generate the low-variance ones, and vice versa. To draw a parallel with the (perhaps more familiar) t-test, if we have two very distant pairs of internally close observations, it is very unlikely that the same normal distribution would generate them all, and we can obtain an arbitrarily low p-value against this null hypothesis as the pair spacing increases (e.g. (0, 0.01) and (0.99, 1) gives p < 0.0001).

### Bonus results II – The case of nonzero selection

Paralleling his work on allele distributions under neutral drift (Kimura, 1955), Kimura also derived distributions for the case of selective pressure favouring one allele (and for the case where this selection fluctuates) (Kimura, 1954). Wright and Kerr (1954) also made quantitative progress on this question; Kimura (1954) links the two approaches. Interpreting these allele frequency distributions as heteroplasmy distributions, we have a probability density function for heteroplasmy measurements given an effective population size and selection coefficient. Once again, a maximum likelihood approach can be used to estimate these parameters and associated uncertainty. Further, a likelihood ratio test can be used to seek statistical support for the presence of selection at the intra-cellular level. In Otten *et al*. (2018), a similar philosophy is used for the inter-cellular level, with a likelihood ratio test against an alternative hypothesis of a “truncated Kimura” distribution – a Kimura distribution with probability mass removed at high heteroplasmies, to model an absence (due to removal or death) of cells with such high heteroplasmies. In both cases, a parametric model for the action of selection (within or between cells) is invoked to provide a well-posed statistical test.

## Discussion

The work of Wonnapinij *et al*. (2008; 2010) was groundbreaking in applying statistical methods and stochastic models from population genetic theory to modern mtDNA observations. The Kimura distribution is a convenient and powerful model for allele frequencies, though as critiqued in Wallace and Chalkia (2013), it is not without issues. Here we suggest that its use within a maximum likelihood setting, rather than using method-of-moments fitting, resolves several issues that have arisen in its application. In particular, we caution against the KS testing of the Kimura distribution fitted by moments. As we have shown, the moment fit does not generally give the parameters that are most compatible (in the KS sense) with the data, making it very likely that false positive errors occur.

This is fundamentally a statistical story about how different estimators can give different results, and how any results must be interpreted with the estimator in mind. There is nothing intrinsically good or bad about the different estimators (method of moments, maximum likelihood, minimum KS distance) that we use here. However, approaches for testing hypotheses with parameters require those parameters to be inferred in a compatible way. When the hypothesis is, for example, any Kimura distribution has a large KS distance from the empirical data, the estimator (minimum KS distance) that minimises this distance and therefore more strictly tests the hypothesis should be chosen. For estimates of uncertainty in sample variance in real datasets, the difference between different estimators rarely provides a substantial effect. However, for fitting and testing distribution structure, the appropriate estimator is a much more important issue.

Wallace and Chalkia (2013) previously outlined some shortcomings of using this approach, noting that the Kimura model itself relies on several assumptions which are not met in real mtDNA situations, and that departures from these assumptions may challenge the model. They also discuss that the use of the KS test to compare the theoretical distribution with empirical data is not without several other issues – including the applicability of its underlying population-genetic assumptions, limited sensitivity, and ability to capture the non-Mendelian dynamics of mtDNA inheritance. The issue we highlight here is a different one and stands in parallel with these important points – even if the Kimura model and KS test approach is accepted, the way that this model is typically fitted with experimental observations makes further testing uninterpretable.

How should we detect selection? In the case of a mechanism that fundamentally changes the family of distributions from which heteroplasmy is drawn, the likelihood ratio test approach of Otten *et al*. (2018) is well suited. Here, support is compared for a truncated Kimura distribution (modelling cell-level selection) and a Kimura distribution (modelling neutrality). The case of intracellular selection is more challenging. Kimura (1954) shows that even in the presence of selection, distributions can closely resemble those from a neutral model. The best approach here is to use longitudinal data (an early reference measurement or a time course) and to fit a model that allows for selection and generates all observations. It is important to note that any early reference measurement will itself be a sample and cannot be regarded as ground truth (i.e. as a population parameter).

The various nonlinearities involved in these expressions and distributions mean that even small differences in individual heteroplasmy measurements -- and certainly in estimated parameters -- can have dramatic differences on the p-values from the analysis (as in Fig. 1B, and several examples in Table 1). Because of this, rounding and binning heteroplasmy values before the WCS-K analysis can have strong effects on the consequent findings (as in some examples in Table 1). The other approaches outlined here are more robust to such small deviations (which will always arise due to measurement noise).

More generally, mtDNA (and ptDNA) heteroplasmy is a remarkably awkward quantity. If we use the near-universal definition *h* = *m* / (*w*+*m*) where *m* is mutant copy number and *w* wildtype copy number, *h* can strictly only take values where *w* and *m* are integers, and where *w*+*m* is cellular copy number. So steps smaller than 0.01 are not permitted for a cell with 100 mtDNAs: we can have *h*=0.05 or *h*=0.06 but not *h*=0.054. As the ratio of two random variables, the variance (and mean) of *h* does not have a convenient closed form representation, even if we have a perfect theory for how *w* and *m* change. The WCS-K model is one attempt to work with *h* by employing a particular set of simplifying assumptions (which have been critically discussed, for example, in Wallace and Chalkia (2013)). Other approaches include attempting to develop predictions for *w* and *m* and then using our approximating a complicated sum over all possibilities (Johnston *et al*., 2015) or a Taylor expansion approximation for *h* (Johnston *et al*., 2015; Hoitzing *et al*. 2019; Edwards *et al*. 2021; Aryaman *et al*. 2019; Insalata *et al*. 2021). These approaches have merit (and shortcomings (Glastad & Johnston 2022)) when attempting to build a bottom-up theory from microscopic dynamics; the Kimura model is convenient for top-down, data-driven analysis. For this reason, it is a valuable approximation of use in heteroplasmy analysis -- providing the estimator used is appropriate for the statistical task in hand.

## Methods

### Statistics and code

The statistical analyses we employ here are described in the text and were implemented in R within a new package *heteroplasmy* (https://github.com/kostasgian21/heteroplasmy). The existing R packages used are: *kimura* (https://github.com/lbozhilova/kimura), *ggplot2* (Wickham, 2016) for plotting, *foreach* (Weston and Microsoft Corporation, 2020) (https://CRAN.R-project.org/package=foreach) and optionally *doParallel* (Weston and Microsoft Corporation, 2022) (https://CRAN.R-project.org/package=doParallel) for parallel execution of the code, and *devtools* (Wickham *et al*., 2020) (https://CRAN.R-project.org/package=devtools) to download and install the *kimura* and *heteroplasmy* packages. The code is freely available at https://github.com/StochasticBiology/heteroplasmy-analysis.

### Heteroplasmy data

The LE and HB mouse oocyte heteroplasmy data were taken from Burgstaller *et al*. (2018). In our paper, only indicative samples of the dataset in Burgstaller *et al*. (2018) were used. LE and HB correspond to two genetically distinct heteroplasmic mouse lines. For the purposes of this work, only samples from oocytes were used. Heteroplasmy measurements were taken as found in the supplementary data file.

Data from progenitor germ cells (PGCs), oocytes, and offspring were taken from Freyer *et al*. (2012). Grouping of individual measurements was undertaken to match the grouping within the original publication, according to the ranges of the mother’s reference heteroplasmy for PGCs and offspring as classified in the paper (i.e., [0.0-0.6],(0.6-0.7],(0.7-1.0]). For the oocyte data, all the 5 samples were individually tested (again, as in the original paper). Because of ambiguity with the order of reporting of the associated significance results in the original paper for the oocyte data, we mapped each of our results to the numerically closest ones from the original paper.

Human, fly (*Drosophila mauritiana* and D. *simulans*), and mouse heteroplasmy measurements from Wonnapinij *et al*. (2008) were also used. Originally, the human data came from Brown *et al*. (2001), fly data from de Stordeur *et al*. (1989), and mouse data from Jenuth *et al*. (1996). The raw data are not available in either publication, so we use the binned data, as depicted in relevant histograms in Wonnapinij *et al*. (2008). To interpret the binned data as estimated individual measurements we tested several methods, including taking the mean of each bin, taking the mean and then adding small noise disturbances to cut the ties, and others. The method that most faithfully reproduced the original results was resampling the data within each bin by taking uniformly random values within the bin interval. As noted also in Wallace and Chalkia (2013), caution is needed when binned data are used to test for selection through fitting a Kimura distribution. Therefore, we reported here ranges of p-values when using the method of moments approach to replicate the results from (Wonnapinij *et al*., 2008), instead of single values (in Table 1).

Finally, organellar heteroplasmy data were taken from Broz *et al*. (2022), including between-generation measurements of mitochondrial heteroplasmy for the *mt91017* and *mt334038* SNPs from MSH1 plants, and the *mt334038* SNP from wild-type plants.

All data, if not presented as such, were normalised to proportions (on the [0,1] interval) rather than percentages for analysis.

For the synthetic data, the normal samples were generated using the *rnorm* command in base R. For the data generated by a Kimura distribution, the *rkimura* function from the *kimura* R package (https://github.com/lbozhilova/kimura) was used.

## Acknowledgements

This project has received funding from the European Research Council (ERC) under the European Union’s Horizon 2020 research and innovation programme (Grant agreement No. 805046 (EvoConBiO) to IGJ). DBS and AKB are supported by NIH Grant R01 GM118046.

## Appendix

### Bottleneck size estimates avoiding the sample mean

Consider the case when we have a known initial heteroplasmy measurement of *h*_0_ = 0.1, then take the previously used observations *<u>h</u>* = (0, 1). Now

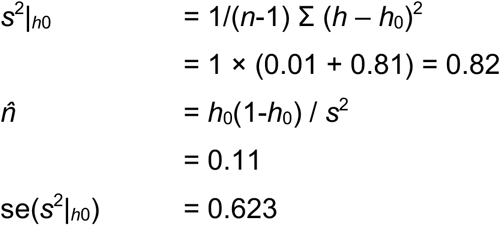

### Numerical issues

An issue can arise when numerically maximising the likelihood associated with homoplasmic data. Because the parameter *b* is constrained to lie on the interval [0,1], we either bound the domain of optimisation or use a transformation to cast the real line onto that interval. In both cases, the gradients in the associated Hessian matrix behave poorly numerically when *b* approaches 0 – as is the case for homoplasmic data, where *b* = 0 is the point estimate. In these cases, we found that using bootstrap resampling to characterise confidence intervals for *b* was the better approach. Of course, bootstrapping could be used for confidence intervals in non-homoplasmic cases too, but for reasons of computational speed and accuracy we suggest Fisher information for estimating *p* in all cases and for estimating *b* in cases of heteroplasmy, and bootstrapping for estimating *b* in cases of homoplasmy.

Another numerical issue involves minimising the KS distance over parameterisations. Because the KS distance surface as a function of parameter *p* and *b* can be non-convex (Fig. 1Aiii), it is hard to guarantee that the best solution has been found: the local optimum found may depend on initial conditions. In our implementation we perform two searches, using the method-of-moments and maximum likelihood parameter estimates as two different initial conditions, and reporting the best solution found from these two searches. We found this to generally identify good solutions based on exhaustively visualising the full surface.

### Kimura fourth central moment

The original article from Kimura (1955) gives an expression for the *n*th moment about zero of the Kimura distribution:

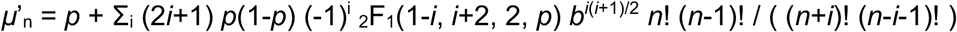

where we have used the *b* = exp(-*t* / *N*_eff_) substitution from Wonnapinij *et al*. (2008).

The infinite sums in this expression terminate when the numerator in the summand contains a factor of zero, hence when *i ≥ n*. The highest *i* value we need to consider is *i* = 4, and for these values the hypergeometric function takes simple polynomial forms: _2_F_1_(1-*i, i*+2, 2, *p*) = {1, 1-2*p*, 1-5*p*+5*p*^2^, 1-9*p*+21*p*^2^-14*p*^3^} respectively for *i* = {1, 2, 3, 4}.

Using these expressions, the non-central moments are then

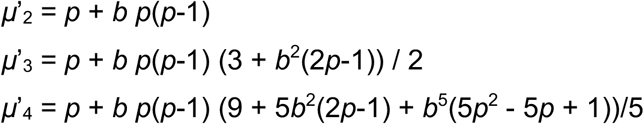

Central moments are related to moments about zero by a family of expressions, of which the fourth is

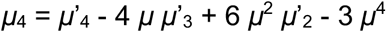

For the fourth central moment we therefore have

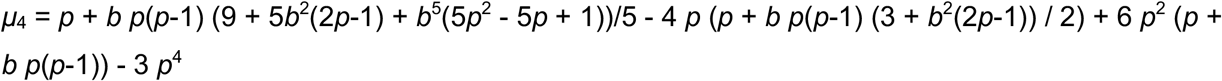

Which is equivalent to the Wonnapinij *et al*. (2010) expression with the identity *σ*^2^ = *p*(1-*p*)(1-*b*).

**Supplementary Figure S1.**
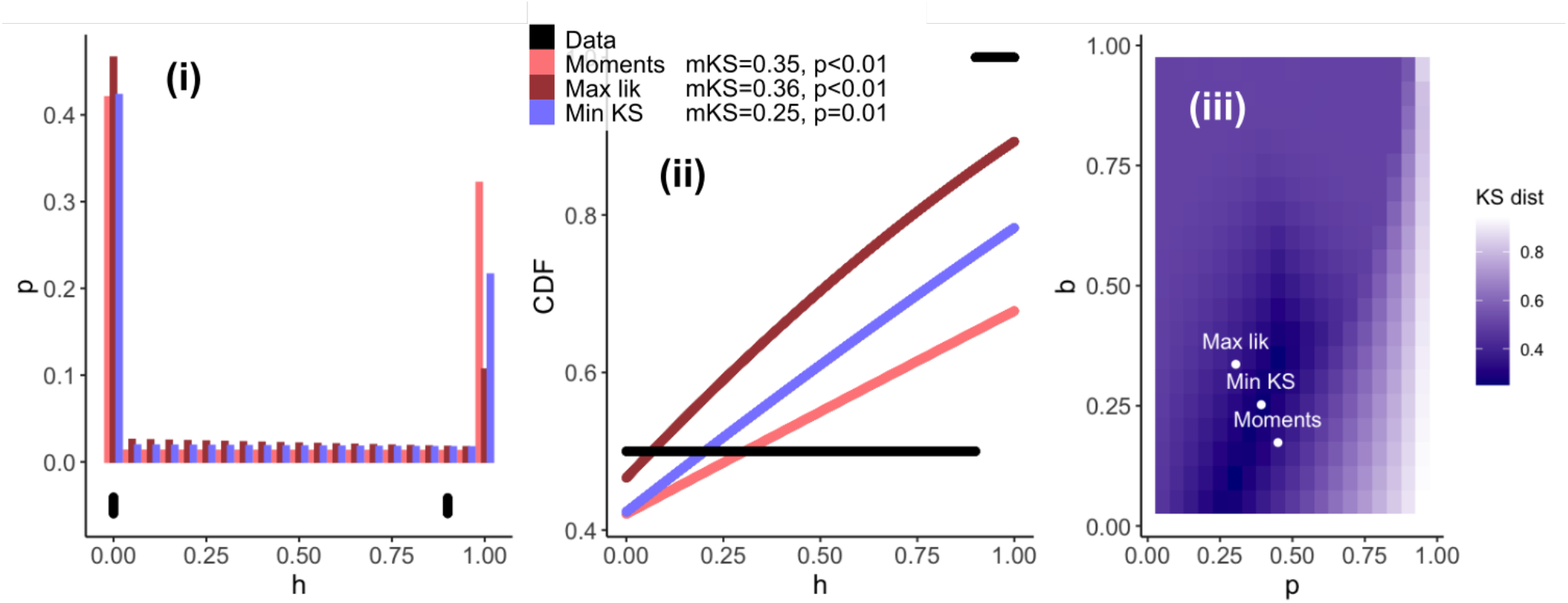
Example of synthetic data that significantly departs from the Kimura distribution. The heteroplasmy measurements consist of the pair (0,0.9) repeated 50 times. As in Fig 1, (i) probability distributions for each fit; (ii) comparison of cumulative distribution functions (CDFs) from the data and for each fit; (iii) the KS distance between empirical observations and theoretical distribution with parameters of the Kimura distribution. mKS, KS distance for each method of fitting; p-values are from the Monte Carlo KS test in WCS-K. As the sample size is higher in this case, the fit using the method of moments and that using maximum likelihood provide more similar parameter estimates.

**Supplementary Figure S2.**
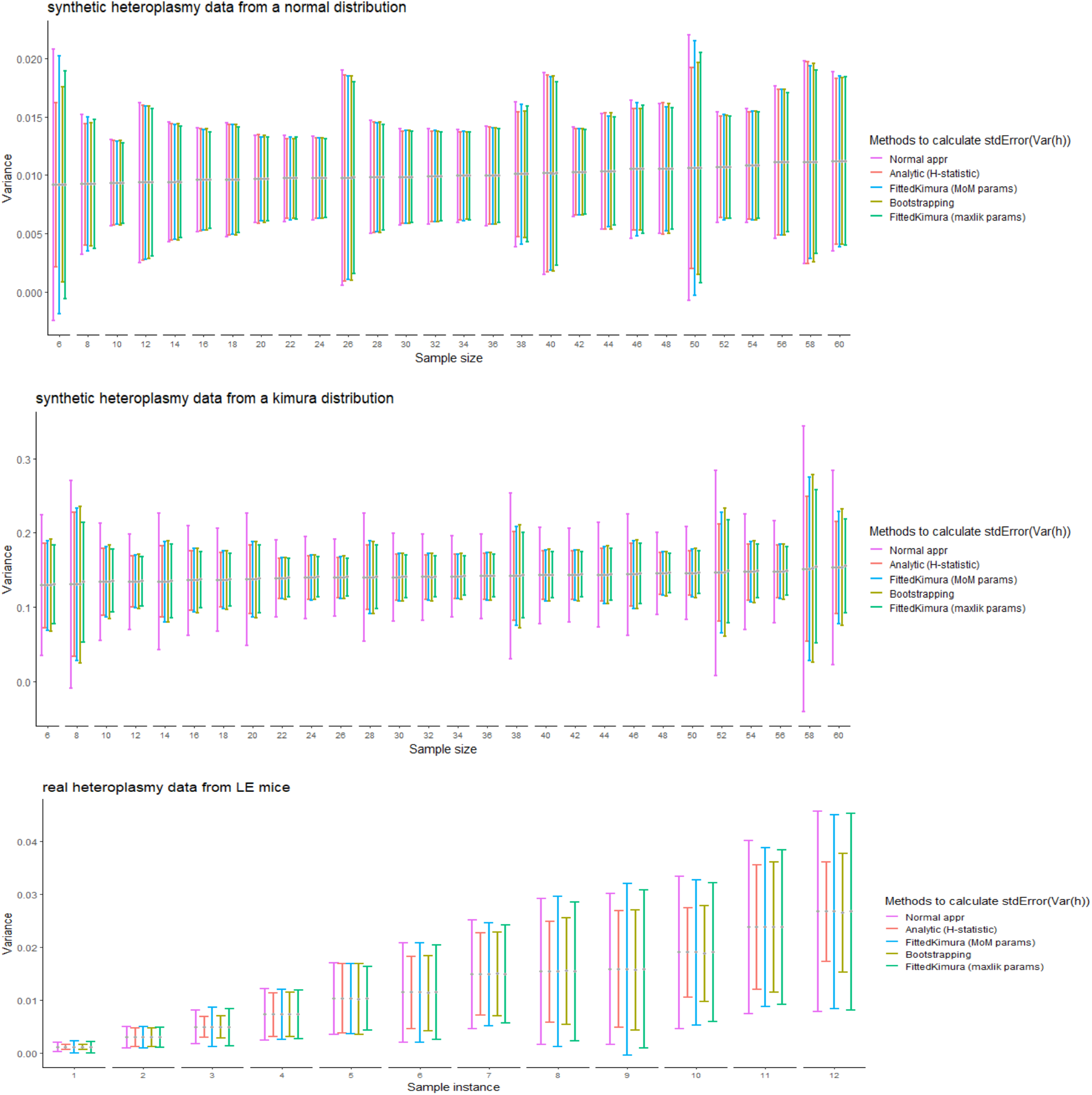
Different ways to estimate the uncertainty of variance in samples from (top) synthetic normal distribution; (centre) synthetic Kimura distribution; (bottom) example mouse data. For the synthetic distributions, samples are ordered by sample size; for the mouse data, 12 example LE type samples are depicted for which the ordering is (for just visual clarity) ascending according to the mean sample variance. Error bars correspond to 2x the standard error of variance as calculated by each method. Analytic (h-statistic) and bootstrapped results are typically comparable; Kimura fits often provide a more conservative estimate of V(h) variance.

